# Rational design of human cathepsin L with efficient collagenolytic activity

**DOI:** 10.1101/2020.05.10.087031

**Authors:** Debi Choudhury, Sampa Biswas

**Author notes:** **Database:** The atomic coordinates and structure factors for CTSL-YLAL and CTSL-C4S mutants of human CTSL (codes 6JD8 and 6JD0 respectively) have been deposited in the Protein Data Bank (http://wwpdb.org/).

## Abstract

Collagens are the main structural components of ECM and collagenolysis is vital for ECM remodelling, which is essential for normal organ homeostasis. Any dysregulation of this process can result in various pathological conditions. The degradation of the interstitial collagens is generally mediated by specific collagenases which are capable of cleaving at specific loci in the collagen triple helix. Lysosomal cysteine cathepsins have general housekeeping as well as some highly specialized functions. Amongst them, human cathepsin K is the only lysosomal cysteine protease which has potent collagenolytic activity against type I collagen. In this study, we have imparted collagenolytic property to another human cysteine protease, cathepsin L, by systematically engineering proline-specificity and GAG-binding surface in the protease. The designed proline-specific mutant shows high specificity for peptidyl substrate containing proline at P2 position but is incapable of cleaving collagen. However, when the proline-specific mutant is further engineered for GAG-binding surface, it can degrade type I collagen in presence of Chondroitin 4-Sulfate (C4-S). We also present high resolution crystal structures of these proline-specific (1.4 Å) and collagen-specific (1.8 Å) mutants. Finally docking studies with prolyl-peptidic substrate (Ala-Gly-Pro-Arg-Ala) at the active site and a GAG molecule (C4-S) at the GAG binding site enables us to identify key structural features responsible for collagenolytic activity of papain-like cysteine proteases. This study provides a rational approach to engineer an efficient collagenase from a human template enzyme which may have various noninvasive therapeutic implications with an expected additional advantage of immuno-compatiblity.

## Introduction

Collagens are essential structural component of the extracellular matrix (ECM) and provide biomechanical properties to connective tissues. This fibrous protein species is the major component of skin and bone and represents approximately 25% of the total dry weight of mammals (1). Different types of collagens are involved in various biochemical and physiological functions. Collagen molecules are comprised of three α chains that assemble together to form a rigid triple helical structure. Every α chain is composed of more than a thousand amino acids based on the repeating sequence Xaa-Yaa-Gly-. The presence of glycine is essential at every third amino acid position in order to allow for a tight packaging of the three α chains in the tropocollagen molecule and the X and Y positions are mostly filled by proline and 4-hydroxyproline, which gives unusual stability of the collagen fibril (2; 3).

Collagenolysis is a physiologically important phenomenon; inadequate collagenolysis leads to accumulation of undigested collagen fibrils in lysosomes within osteoclasts (4), as observed in patients with the autosomal recessive skeletal dysplasia pycnodysostosis (5) and on the other hand, uncontrolled collagenolysis leads to different pathological conditions (6) such as rheumatoid arthritis (7), atherosclerosis (8), pulmonary emphysema (9), tumor invasion and metastasis (10). Therefore critical understanding of collagen degrading enzymes or collagenases, their function, spatio-temporal expression, regulation of their proteolytic activity and their inhibition, is important. In addition to medical and pharmaceutical industry, collagenases have been used in food, cosmetics and textiles segments and have applications in fur and hide tanning to help ensure the uniform dying of leathers (11; 12). In medical applications, it can be used in burns and ulcers treatment (13;14), to eliminate scars, (15), for Dupuytren’s disease treatment and to various types of fibrosis such as liver cirrhosis, for preparing samples for diagnosis, for production of peptides with antioxidant and antimicrobial activities (16) and play an extremely important role in the transplant surgery success of some specific organs (15). Thus, an efficient collagenase will definitely have immense utilization in various pharmacological and commercial applications.

The ECM, being a highly dynamic structure, constantly undergoes remodeling processes. ECM dynamics are indispensible during restructuring of tissue architecture and tightly controlled ECM homeostasis is crucial for regulating many essential cellular processes, allowing for correct organism development, wound healing and normal tissue homeostasis. Collagens are continually degraded and re-synthesized during the remodelling process of ECM. The complex triple helical fold renders collagen extremely resistant to proteolytic attack and only a few classes of proteases are able to degrade this molecule; among them, the most prominent members belong to the matrix metalloproteinases (MMPs) family e.g. MMP1, MMP8, and MMP13 and some lysosomal cysteine-cathepsins, like Cathepsin B, L and K. The MMPs and Cathepsin K are able to hydrolyze peptide bonds in the triple helical domain of these collagens. The matrix metalloproteinases are a family of Ca^2+^ and Zn^2+^-dependent neutral endopeptidases collectively capable of degrading essentially all components of the ECM. MMPs employ a specific structural element, called the hemopexin domain, to partially unfold the triple helix and cleave at a single site (17; 18). Cathepsins do not possess such a specific “unwinding” domain and most of them are only capable of cleaving in the nonhelical telopeptide regions of collagens (19; 20). The only exception is cathepsin K which can cleave peptide bonds at multiple sites within the triple helical domain (21; 22).

This unique collagenolytic property of CTSK has two major aspects: (i) its proline-specificity i.e. ability to accommodate proline in its active site since collagen fibers are mainly made up of prolines and hydroxyprolines; and (ii) its GAG-binding property i.e. formation of an oligomeric complex between cathepsin K molecules and extracellular matrix-resident glycosaminoglycans (GAGs) which assists its collagenolytic activity by allosteric modulation (23).

There is growing evidence that, in addition to Cathepsin K, other cysteine proteinases also participate in osteoclast-mediated proteolytic degradation of organic bone matrix (24;25;6; 26). In particular, Cathepsin L (CTSL), a lysosomal endo-peptidase mainly involved in tissue degradation and ECM remodeling, appears to make a significant contribution to this process (24; 6; 26; 27). CTSL cleaves only degraded/unfolded collagens like gelatin (28). It is a ubiquitously expressed major representative of the papain-like family of cysteine proteinases. It is also the most active of the lysosomal cysteine proteases with regard to its ability to hydrolyze general protein substrates (29) and is more active proteolytically than CTSK.

We have attempted to generate a mutant recombinant collagenase which is able to cleave the collagen fibers in a similar manner as CTSK. For this, we selected recombinant human Cathepsin L as the target protease. This is because CTSK and CTSL have some functional redundancy and similar catalytic domains and 3D structures (30) and may also have greater antigenic compatibility for pharmacological usage as compared to the microbial or fungal collagenases which are being exploited for such purposes.

The primary specificity of papain-like proteinases is largely determined by the S2-P2 subsite interaction. Contrary to cathepsin L, which accommodates hydrophobic aromatic residues in its S2 subsite, analyses of cleavage sites of cathepsin K in type I and II collagens as well as profiling of its substrate specificity using positional scanning peptide libraries have confirmed that cathepsin K has a more restricted pocket and exhibits an exclusive preference for Pro at P2 among mammalian cathepsins (21). Hence, to be able to degrade intact collagen, an engineered collagenase must be able to accommodate Proline in its P2 position. In this study, we have generated a Proline-specific CTSL by mutating relevant residues in the active site of CTSL as a first step towards producing a recombinant collagenase.

Early experiments suggested that complex formation with Chondroitin Sulfate (CS) is necessary for the collagenolytic activity of cathepsin K (31) though a recent finding has shown that type I collagen can also be degraded even in absence of glucosaminoglycans (32). Nonetheless, bone-resident GAGs have been shown to potentiate the collagenolytic activity of cathepsin K and endogenous GAG concentrations in bone were sufficient for a maximal effect on cathepsin K activity (32). The crystal structure of cathepsin K-Chondroitin 4-Sulfate (C4-S) complex revealed the structural basis for the interaction (33), the binding site being located on the R-domain of cathepsin K interactingwith three disaccharide units of C4-S. The interaction is stabilised mostly by electrostatic potential between the negatively charged C4-S and positively charged residues on the protease surface. To generate the GAG-binding site, we have substituted relevant residues on the proline-specific mutant of CTSL by C4-S interacting residues of CTSK surface to mimic an equivalent positively charged electrostatic surface on CTSL mutant.

Our results show that the structure-based rationally designed, proline-specific mutant of CTSL generated in this study gains proline-specificity for peptide substrate having proline at P2 position and the mutant having both, proline-specificity as well as GAG-binding surface shows distinct collagen degrading activity against triple helical type I collagen. This study further illustrates that just proline-specificity is not enough for cleaving triple helical collagen; there is a requirement for GAG (like chondroitin 4-sulfate here) binding property to efficiently cleave collagen. Therefore from this study, importance of GAG binding for collagen degradation is further attributed for cysteine cathepsins. The crystal structures of the Proline-specific CTSL mutant (CTSL-YLAL mutant) and the proline-specific mutant with GAG-binding site (CTSL-C4S mutant) have been solved at 1.4Å and 1.8Å respectively. The structural analyses along with docking studies with proline containing polypeptide substrate at catalytic cleft and C4-S at GAG binding surface correlates the molecular basis of engineered proline specificity and collagenase activity of cathepsin L mutants.

## Results and Discussion

### Rational design and generation of the proline-specific mutant CTSL-YLAL of CTSL

As is known the primary specificity of papain-like cysteine proteinases is largely determined by S2-P2 interactions. Gly-Pro-X is a repetitive motif in triple helical collagens and it has been demonstrated earlier that CTSK exhibits a preference for proline residues in the P2 position of the substrate (22; 34). The accommodation of a P2 proline residue in the S2 subsite pocket [subsite nomenclature from Schechter and Berger (35) is unique for CTSK and in contrast to the preferred acceptance of bulky aromatic P2 residues by CTSL (Phe>>Leu). We compared the three-dimensional structures of cathepsins K and L [pdb id: 3C9E; pdb id: 1CJL] (36) which showed amino acid sequence differences at residues 69, 161, 164 and 214 in the primary S2 binding pocket (CTSL numbering is used in this paper). The S2 binding pocket residues in CTSL, CTSK and the designed mutation to generate proline specific mutant CTSL-YLAL are shown in Table 1.

**Table 1:**
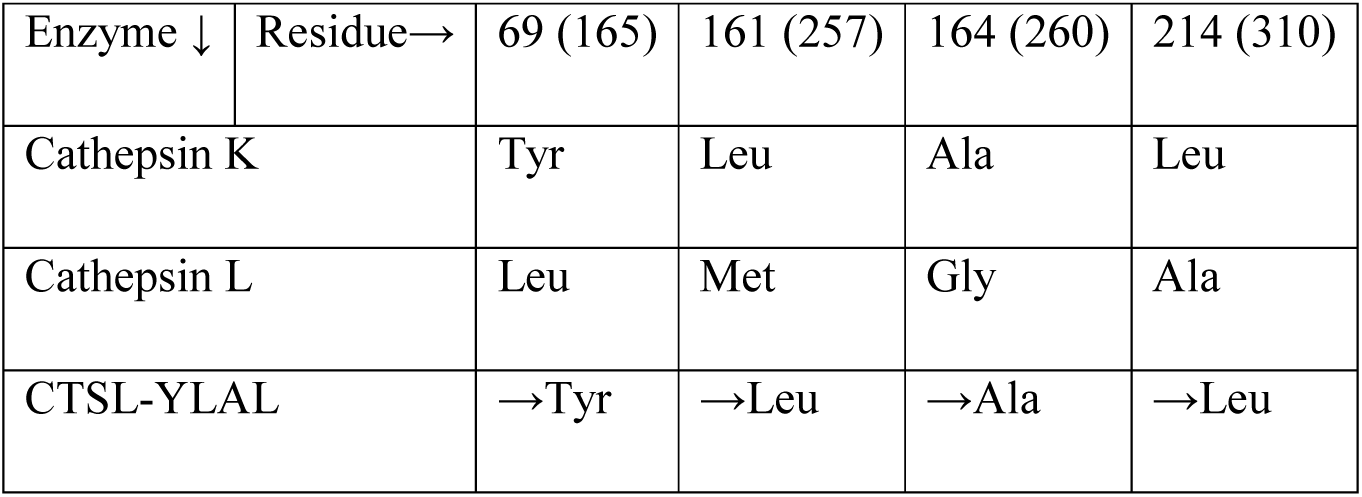
S2 subsite constituting residues in CTSK, CTSL and in proline-specific quadruple mutant of CTSL (CTSL-YLAL) (numbers in brackets are pro-CTSL residue numbers)

Analyses of 3-D structures of the S2 pockets in cathepsins K and L revealed significant differences in size and shape of the respective subsites. Residue 214 (310), located at the bottom of the pocket, and residues 69 (165), 135 (231), and 161 (257) lining the wall of the S2 pocket, tend to determine whether the pocket is larger or smaller (Fig 1A, 1B, 1C and 1D). The S2 pocket of CTSK is more constricted than that of CTSL because of the difference in the amino acid composition (Table 1) of this subsite making it ideal for accommodation of amino acids with smaller side chains like proline and leucine at P2 position of the substrate, whereas the S2 subsite of CTSL can accept bulkier aromatic side chains like that of phenylalanine. It has been shown earlier that whereas proline specificity of CTSK was considerably compromised when a CTSL-like Tyr69Leu/Leu214Ala mutant of CTSK was generated; there was an enhancement of preference for proline in CTSK-like CTSL Leu69Tyr/Ala214Leu mutant (37). Apart from these two residues, we wanted to look at the contribution of the other residues of the S2 pocket also towards the proline specificity of the enzyme and hence we targeted all the 4 residues lining the S2 subsite of CTSL mutating them like that in CTSK and thus generating a quadruple mutant, L69Y/M161L/G164A/A214L (CTSL-YLAL) of cathepsin L. This was also the first step towards producing a mutant CTSL with collagenolytic property.

**Figure 1:**
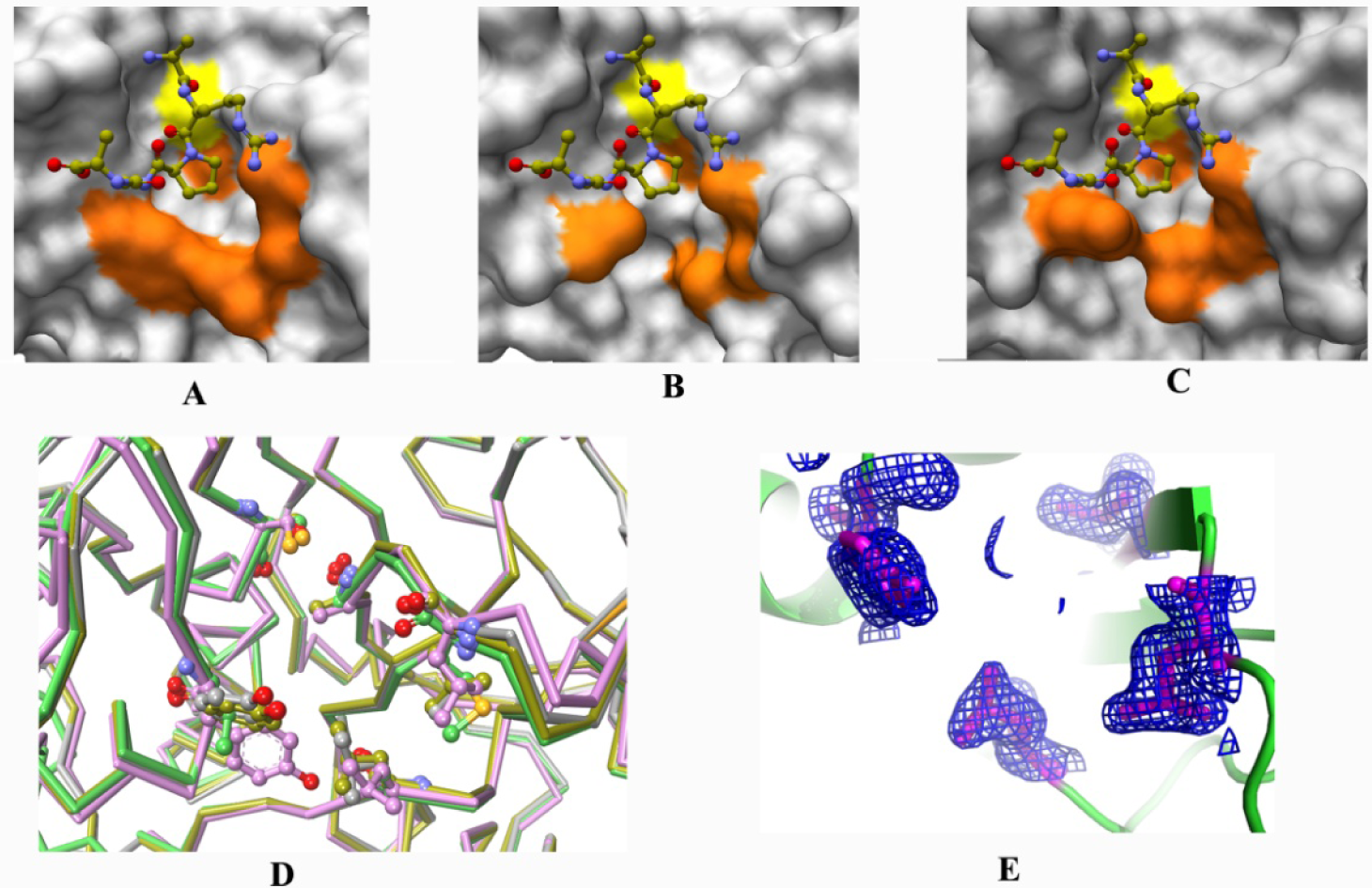
Comparison of S2 pockets. **A**. wild-type Cathepsin K (pdb_ids 3C9E), **B**. wild-type Cathepsin L (pdb_ids 1CJL), **C**. and proline-specific Cathepsin L mutant, CTSL-YLAL, with a docked substrate analog penta-peptide substrate (Ala-Gly-Pro-Arg-Ala). Orange color represents the residues lining the S2 pocket considered for proline specific mutation (Table 1). Yellow color represents the catalytic cysteine, **D**. Superposition of the S2 pocket amino acid side-chain conformations of the above 3 (magenta, green, grey ribbon respectively) proteins and C4S mutant (yellow ribbon), **E**. electron density (at 1Σ) of the mutated residues of S2 pocket in CTSL-YLAL mutant.

The said residues were mutated using appropriate synthetic mutagenic oligo nucleotides and confirmed as mentioned in Materials and Methods section. The pro-form of the mutant was expressed in E. coli as inclusion bodies (Mr 43000) using the pET30/EK LIC vector, purified, refolded using the same protocol like wild-type (Fig 2) and processed to mature form at pH 4.0 in presence of 2 mM EDTA and 20 mM Cysteine and incubation at 45 °C for 10 min.

**Figure 2:**
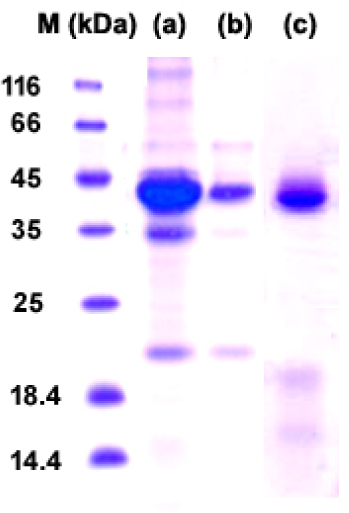
SDS-PAGE analysis of purified pro form of wt-Cathepsin L and its mutants. Lanes (a) wt-CTSL, (b) CTSL-YLAL and (c) CTSL-C4S.

### Enzymatic characterization of recombinant wild type CTSL and CTSL-YLAL mutant using chromogenic substrates

To study whether the mutation of the four residues (Leu69, Met161, Gly164 and Ala214) of the S2 pocket of cathepsin L to that like cathepsin K S2 pocket (Tyr69, Leu161, Ala164 and Leu214) induce expected alteration in the substrate specificity, hydrolysis of three representative chromogenic tripeptide substrates (pEFL↓pNA, VLK↓pNA, GPR↓pNA) with Phe, Leu and Pro at the P2 position respectively, were evaluated and compared with that of wild type CTSL. Steady-state kinetics was performed as described in Materials and Methods, and the experimental data are summarized in Table 2.

**Table 2.**
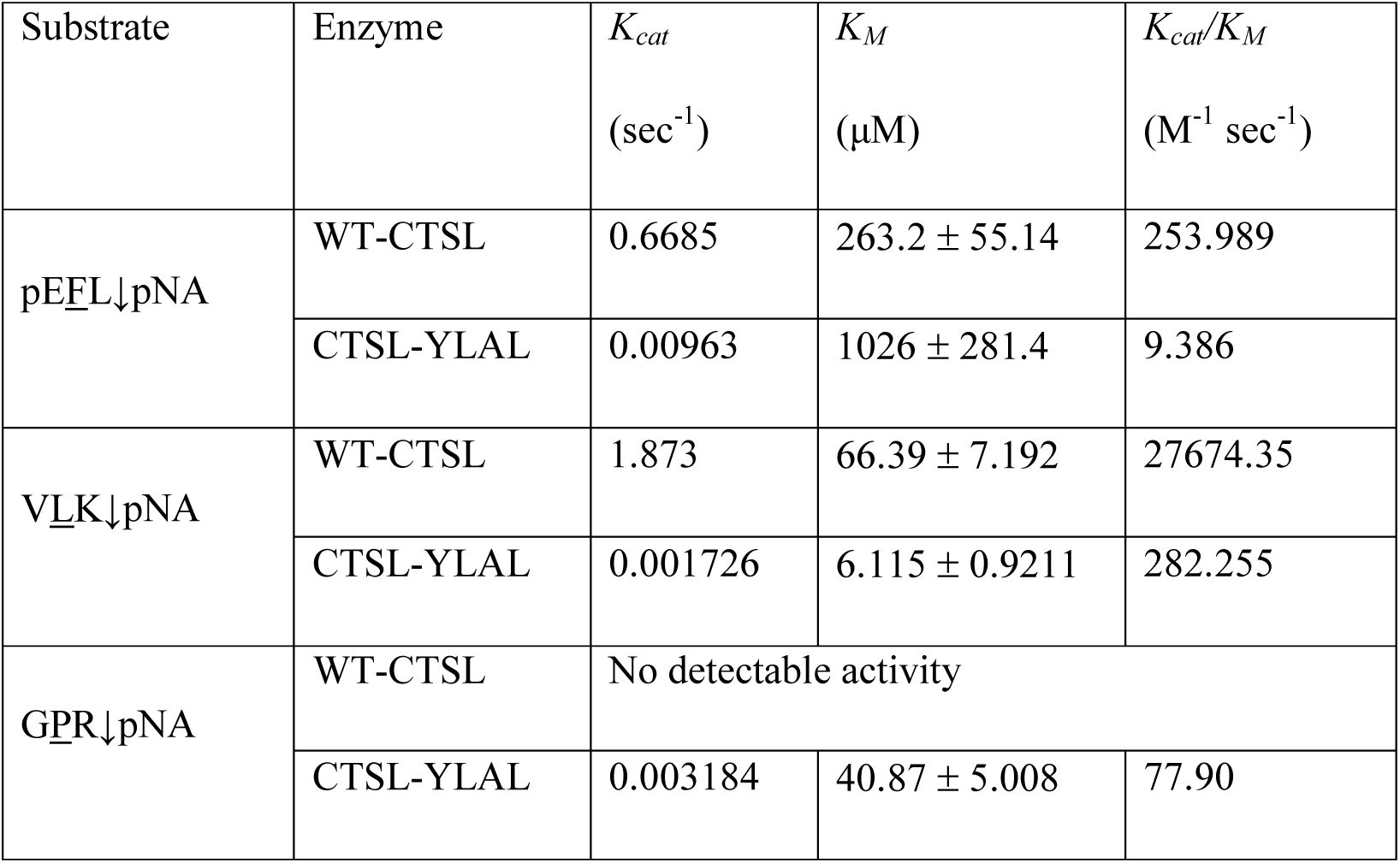
Hydrolysis of peptidyl-pNA substrates by wild-type CTSL and proline-specific mutant CTSL-YLAL

| Substrate | Enzyme | $K_{cat}$<br>(sec<sup>-1</sup>) | $K_M$<br>(μM) | $K_{cat}/K_M$<br>(M<sup>-1</sup> sec<sup>-1</sup>) |
| --- | --- | --- | --- | --- |
| pEFL↓pNA | WT-CTSL | 0.6685 | 263.2 ± 55.14 | 253.989 |
|  | CTSL-YLAL | 0.00963 | 1026 ± 281.4 | 9.386 |
| VLK↓pNA | WT-CTSL | 1.873 | 66.39 ± 7.192 | 27674.35 |
|  | CTSL-YLAL | 0.001726 | 6.115 ± 0.9211 | 282.255 |
| GPR↓pNA | WT-CTSL | No detectable activity |  |  |
|  | CTSL-YLAL | 0.003184 | 40.87 ± 5.008 | 77.90 |

Since our aim was to generate a proline-specific CTSL, we compared the kinetic parameters of the mutant CTSL-YLAL with wild-type CTSL. As can be observed from the kinetic parameters (Table 2), wild-type cathepsin L exhibits a significant preference for Leu>>Phe at P2 position. On the other hand, proline in the P2 position is well accepted by the CTSL-YLAL mutant. There is significant decrease in specificity for Phe at P2 position of the CTSL-YLAL mutant compared to that of the wild-type CTSL indicating a change in the shape and size of the S2 pocket due to replacement of some of the residues which was subsequently supported by our structural studies. Importantly, the kinetic data with G<u>P</u>R↓pNA indicates that the CTSL-YLAL mutant is able to accept Pro at the P2 position because of the introduced mutations in the S2 pocket, whereas the wild-type protein does not show any detectable activity with this substrate.

Earlier studies on a CTSK-like double mutant [L69Y/A214L (L67Y/A205L)] of cathepsin L showed that the specificity (*k*_cat_/*K*_M_ values) decreased by five times for fluorogenic peptide substrate having Phe at P2 position, while there was a 30 times enhancement of the specificity for substrate having Leu at P2 position (37). Their studies also showed that there was a ten times gain of specificity for a Proline containing substrate. Our result with quadruple mutant (L69Y/M161L/G164A/A214L) of CTSL showed loss of specificity for both Phe and Leu containing substrates while significant gain of specificity for Pro containing substrate. This we presume is because of the combined effect of the M161L and G164A mutations apart from the L69Y and A214L mutations, which has further altered the characteristic of the S2 subsite for more stringent specificity for a proline residue of the substrate.

High resolution crystal structure of CTSL-YLAL mutant shows a constriction at the lower part of the S2 pocket has been engineered like CTSK (Fig 1A). A docking study was performed with a penta-peptide substrate Ala(P4)-Gly(P3)-Pro(P2)-Arg(P1)-Ala(P1’) having same residues at P1, P2 and P3 positions of the substrate (G<u>P</u>R↓pNA) used for kinetic experiments. This docking study indicate that, the generated constriction in S2 subsite of the mutant can provide a better interaction / shape complementary to the proline residue of the substrate compared to its wild-type counterpart (Fig 1A, 1B and 1C). The S2 pocket of CTSL-YLAL almost mimics CTSK (Fig. 1A and 1C). However rotamer conformations of the mutated residues L69(165)Y and A214(310)L are slightly different than those of CTSK (Fig 1D and 1E). In the structure of CTSL-C4S mutant, these rotamer conformations of L69(165)Y and A214(310)L are preserved (Fig. 1D) which corroborates the experimental data of the mutant having proline specificity described in the next section.

### Generation of GAG-binding surface on Proline-Specific CTSL and Degradation of triple-helical Type I collagen by collagenolytic CTSL mutant

Cathepsin K forms collagenolytically active complexes with GAGs, with different GAGs competing for the binding to the enzyme (38; 33). Some of them, such as chondroitin and keratin sulfate, enhance the collagenolytic activity of cathepsin K, whereas heparin sulfate or heparin selectively inhibits this activity. CTSL is also known to have extracellular matrix proteins degrading properties (39; 40) and some collagenolytic property. But as mentioned earlier, it can cleave collagen only in their non-helical telopeptide regions, unlike CTSK which can cleave the intact triple helical collagen structure.

The crystal structure of the complex of wt-CTSK and chondroitin-4-sulphate revealed that 12 amino acid residues from R-domain of cathepsin K interact directly or, indirectly via water molecules with the negatively charged C4-S (33). Of these 12 residues, nine amino acid residues interact directly with the sugar (Table 3). It has been shown earlier by Cherney et al. (41) that by replacing the equivalent C4-S interacting residues in CTSK with the appropriate amino acid residues present in CTSL, the collagenolytic activity of CTSK could be altered. Among the various mutants generated in that study, the M5 variant (K9E/I172E/Q173S/N195M/K196G/L200K, CTSL numbering), comprising six substitutions showed the lowest collagenase activity with a reduction of 60% when compared with the WT enzyme. On the other hand, the M3 variant (N195M/K196G/L200K) was an even better collagenase than the wild type peptidase probably because of additional interactions generated between Lys200 and C4-S. The M4 variant (K9E/N195M/K196G/L200K) was found to be significantly more efficient than variant M1 (K9E), for which they postulated that the inhibitory effect of K9E substitution was compensated by the Lys200 interaction in this mutant. In our study, we targeted the direct C4-S interacting residues of CTSK and replaced three of them (Glu9, Met195 and Gly196) with those like in CTSK (Lys9, Asn195 and Lys196) (Table 3) presuming that these three mutations may induce C4-S binding property to CTSL. All the three substituted residues are positively charged which is likely to assist the binding of the negatively charged C4-S molecule to the peptidase. For this reason, the residue Lys200 of CTSL was not substituted since it already has positive charge. The mutations were performed on the proline-specific CTSL, CTSL-YLAL, confirmed by sequencing, expressed and purified as mentioned in Materials section and named as CTSL-C4S mutant.

**Table 3:** Residues interacting directly with C4-S molecule in CTSK-C4-S complex structure (pdb id: 3C9E), residues in equivalent position in CTSL and designed mutations of CTSL for C4S binding.

| Residue* | Cathepsin K | Cathepsin L | CTSL-C4S mutant |
| --- | --- | --- | --- |
| 6 (102) | Asp | Asp | - |
| 8 (104) | Arg | Arg | - |
| 9 (105) | Lys | Glu | →Lys |
| 10 (106) | Lys | Lys | - |
| 172 (268) | Ile | Phe | - |
| 173 (269) | Gln | Glu | - |
| 195 (291) | Asn | Met | →Asn |
| 196 (292) | Lys | Gly | →Lys |

\*Mature CTSL numbers has been used with Pro-CTSL numbers in brackets

Type I collagen from calf skin was incubated with free wild-type CTSL; proline-specific mutant, CTSL-YLAL; and the collagenolytic mutant, CTSL-C4S, for 2h, 4h and overnight at 28 °C and for 2h and 4h at 37 °C in presence and absence of chondroitin 4-sulfate (Fig 3A, 3B, 3C and 3D). At 28 °C, in the presence C4-S, enrichment of α-bands was observed at 2 h and 4 h of incubation with gradual disappearance of β- and γ-bands (Fig 3A and 3B). Thus there was increase in the degradation of type I collagen by the CTSL-C4S mutant protease in presence of C4-S with increase in incubation period and the collagen was almost completely degraded after overnight incubation at 28 °C (Fig 3B). Incubation at 37 °C in presence of C4-S showed significant collagen degradation after 2 h incubation, and there was total disappearance of all the collagen bands after 4 hours of incubation almost like recombinant human cathepsin K in a similar experiment from our laboratory (42). On the other hand, the wild-type CTSL and the CTSL-YLAL mutant showed no degradation of the extracellular matrix protein even after overnight incubation either at 28 °C or at 37 °C (Fig 3A, 3C and 3D).

**Figure 3:**
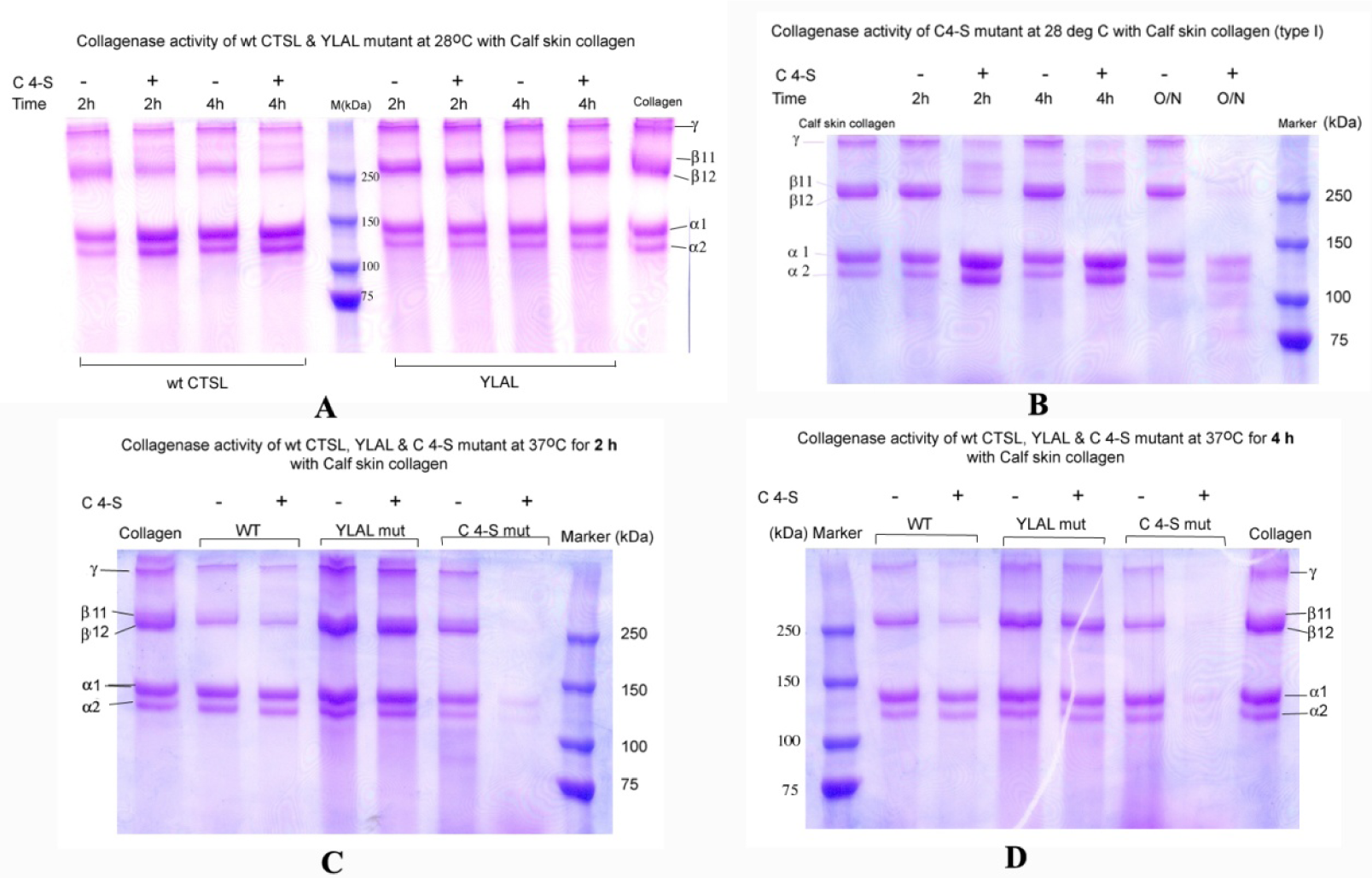
Type I calf skin collagen degradation by wt cathepsin L, proline-specific cathepsin L mutant (CTSL-YLAL) and collagenolytic cathepsin L mutant (CTSL-C4S) in presence and absence of chondroitin 4-sulfate. **A**. wt CTSL and CTSL-YLAL at 28 °C for 2 h and 4 h, **B**. CTSL-C4S at 28 °C for 2 h and 4 h, **C**. wt CTSL, CTSL-YLAL and CTSL-C4S at 37 °C for 2 h, **D**. wt CTSL, CTSL-YLAL and CTSL-C4S at 37 °C for 4 h.

Chondroitin 4-sulfate is a highly negatively charged molecule requiring a positive electrostatic surface for appropriate binding which is aptly provided by CTSK. We replaced three residues (Glu9Lys/Met195Asn/Gly196Lys) in the corresponding surface region of CTSL-YLAL mutant so as to mimic the CTSK surface. The amino acid replacements, Glu9Lys/Met195Asn/Gly196Lys, resulted in a positively charged surface appropriate enough for binding the electronegative C4-S molecule so as to aid the collagenolytic activity of the CTSL-C4S mutant (CTSL-YLAL + C4-S/GAG binding region) as is apparent from the collagen degradation experiments as well as the electrostatic surface diagram of both the proteases (FIG 4A, 4B and 4C). In an earlier study, it has been shown that a combination of six amino acid replacements (K9E/I171E/Q172S/N190M/K191G/L195K) in CTSK with those of CTSL displayed a cumulative effect and inhibited the collagenolytic activity (41). Thus, we present here experimental evidence that replacement of only three amino acids can induce chondroitin 4-sulfate binding and subsequent collagenolytic property in a non-collagenolytic cysteine protease, cathepsin L.

**Figure 4:**
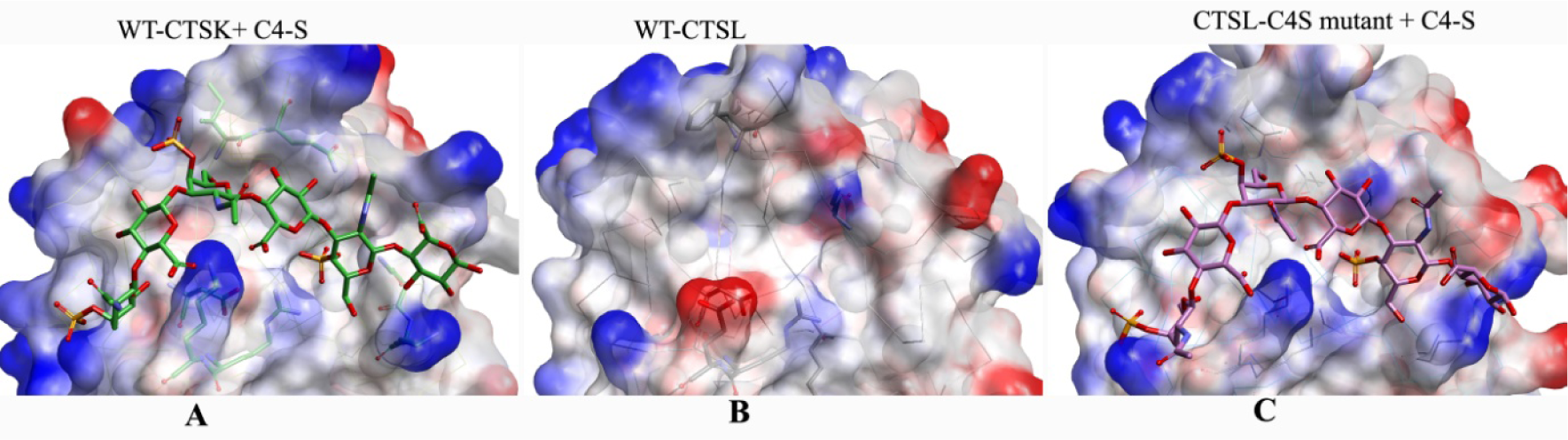
Surface representation of wt CTSK, wt CTSL and CTSL-C4S mutant showing change in electrostatic property of the CTSL GAG-binding region. A. CTSK with attached chondroitin 4-sulfate (pdb id: 3C9E), B. the equivalent surface of wild type CTSL (pdb id: 1CJL) and C. the CTSL-C4S mutant of CTSL as result of 3 mutations (E9K/M195N/G/196K) replacement with a C4-S hexasaccharide docked in the binding groove.

Engineered GAG binding surface of the CTSL-C4S mutant is occupied by water and organic solvent molecules in the crystal structure (Fig 5A and 5B) almost covering the C4-S binding region found in crystal structure CTSK-C4-S complex (Fig 5C). We also docked a C4-S molecule by replacing these ligands in the newly generated surface of CTSL-C4S mutant and the complex structure shows that the mutations provide correct electrostatic surface for C4-S binding (Fig 4C).

**Figure 5:**
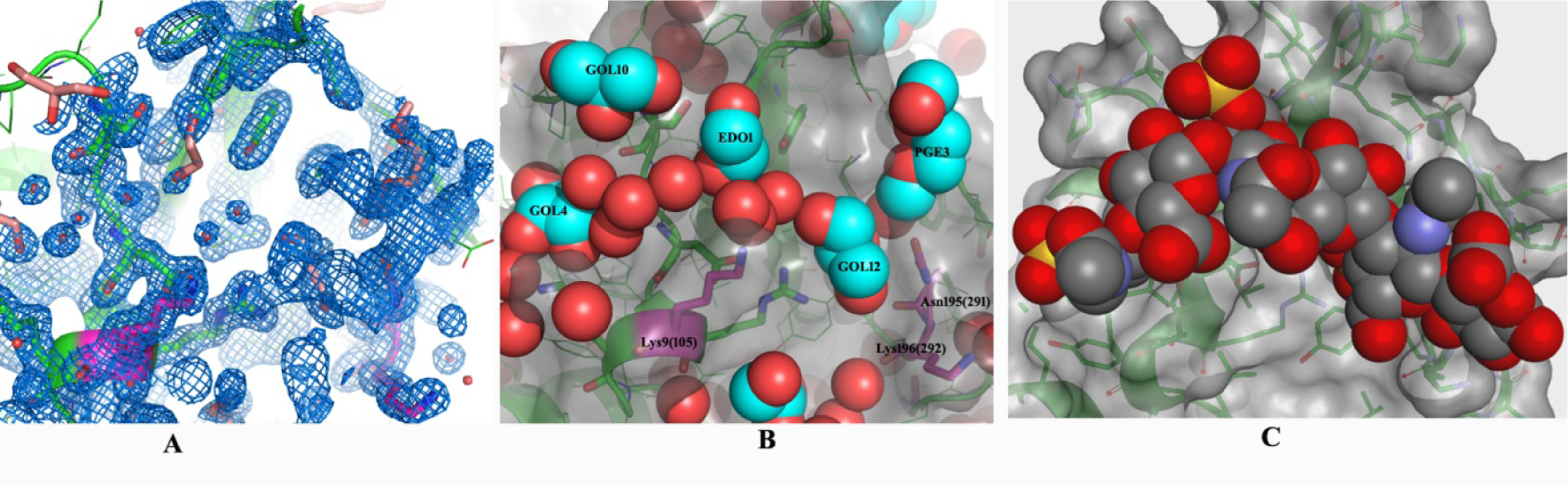
Chondroitin 4-sulfate binding region in the crystal structure (pdb_id: 6JD0). **A**. Electron density at 1Σ. **B**. The organic solvent ligands in the crystal structure occupying the GAG binding surface in Van der waal presentation. The mutated residues are shown in magenta color. **C**. Equivalent GAG binding region of CTSK with C4-S (pdb id: 3C9E).

Analyses of the interaction between the protein and the C4-S molecule in the structure revealed that Glu9 (105) to Lys mutation in CTSL-C4S mutant provides the main anchoring interactions of the middle part of C4-S polymer while Met195 (291) to Asn and Gly196 (292) to Lys mutations provide proper electrostatic environment to stabilize the end (Fig. 6A, 6B and 6C). Superposition of C4-S chains in CTSK-C4-S complex structure (pdb_id: 3C9E) and that in docked conformation with CTSL-C4S mutant, shows a similar curvature of the chains with slight orientation difference in the sugar moiety of the first glucuronic acid residue BDP301. This alteration of orientation is mainly due to difference in electrostatic environment, still exists in the mutant structure (upper part of BDP301) compared to CTSK (Fig. 6A, 6B and 6C).

**Figure 6:**
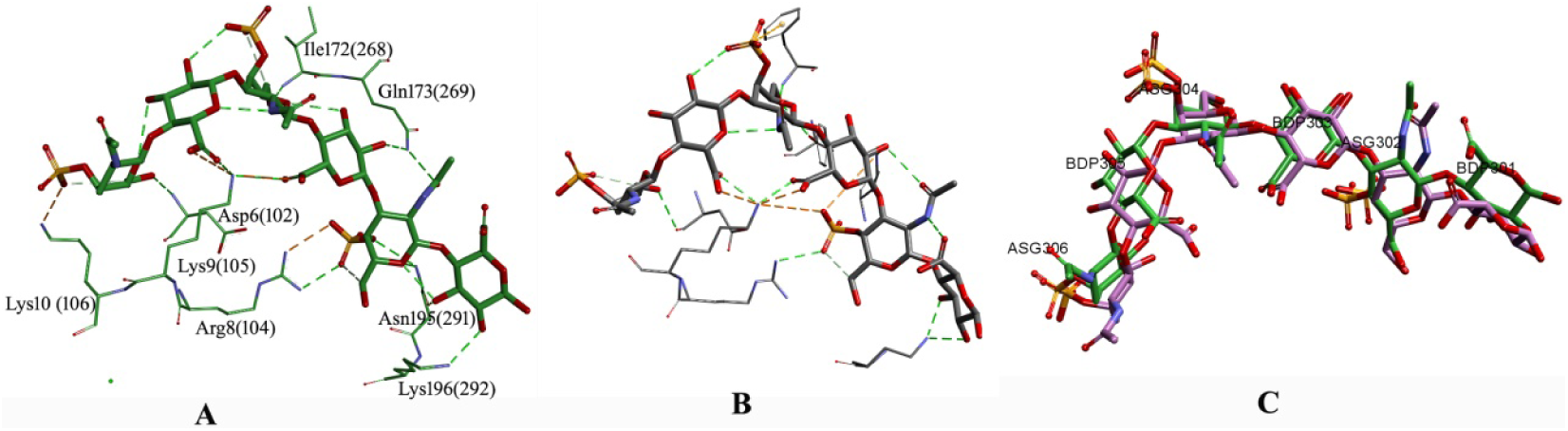
C4-S-protein interactions between the C4-S hexasaccharide and CTSK (pdb id: 3C9E) **A**., between C4-S and CTSL- C4S mutant **B**. and **C**. superposition of C4-S molecules in CTSK and CTSL-C4S mutant. The β-D-glucuronic acid (GCU1) is labeled, as is the 2′-deoxy-2′-acetamido-β-D-galactose-4-sulfate (ASG6). Amino acid labelling id is CTSL based as given in Table 3.

### Proline specificity and influence of GAG in general proteolytic activity of CTSL-C4S mutant

Kinetic analyses with proline containing substrate G<u>P</u>R↓pNA reveals that CTSL-C4S mutant retains its proline specificity almost like CTSL-YLAL mutant having *K*_*M*_ values in the same order. It has also been observed that GAGs, C4-S and heparin (HP), significantly enhance general proteolytic activity of the CTSL-C4S mutant against the same substrate.

Earlier it was reported that (43) heparin exerts different effects on cathepsin K in terms of activity and stability despite similarity with other GAGs. Such strong effects of heparin on CTSL-C4S mutant have also been observed in our study with much higher activity (Fig. 7A) and formation of an oligomeric state of the CTSL-C4S mutant as revealed in Native-PAGE (Fig.7B). This effect of HP can be attributed by its high negative charges compared to other GAGs. However our observations clearly indicate engineering of GAG binding positively charged surface in human CTSL (Fig.7C) has a similar GAG binding effect as it is observed in CTSK.

**Figure 7.**
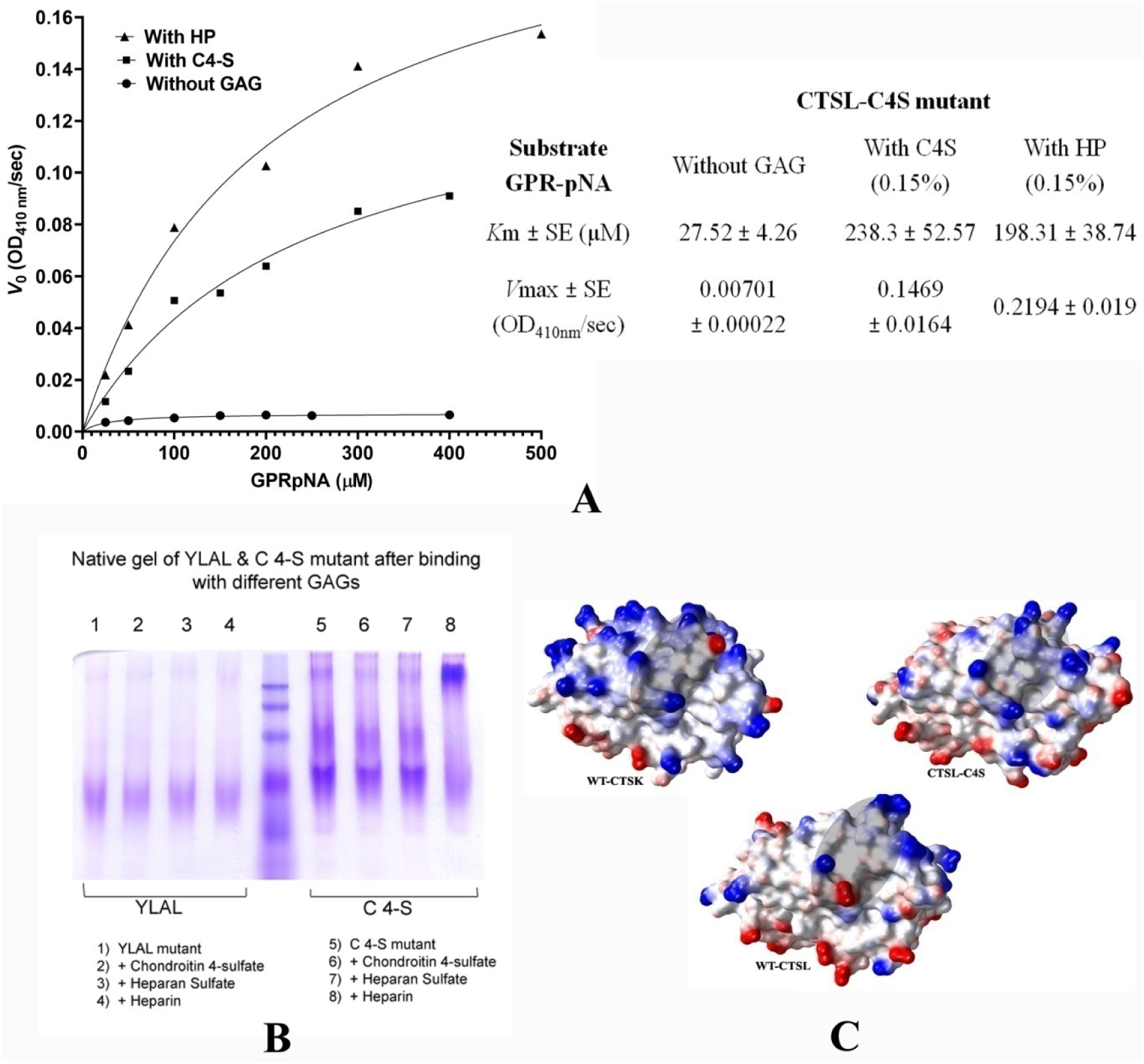
Effect of GAG in CTSL-C4S mutant protein. **A**. Michaelis plot and corresponding kinetic data of CTSL-C4S mutant in presence and absence of GAGs. **B**. Native PAGE of CTSL-YLAL and CTSL-C4S mutants in presence and absence of different GAGs. **C**. Change in electrostatic surface potential in CTSL-C4S mutant; a comparison with wild-type CTSL and CTSK. Significant changes are shaded.

## Conclusion

Crystal structures of CTSL-YLAL and CTSL-CS4 mutants are in the pro-form with active site Cys mutated to Ser to avoid auto catalytic degradation. Our objective was to understand the effect of mutations in the structure of entire zymogenic form of the protease. Though both the mutants refolded, purified and crystallized in a similar condition, the crystals are not isomorphs with noticeable difference in their cell dimensions (Table 5), Vm values (1.80 and 2.32 respectively) and solvent content (31.7% and 47.0% respectively). Both the crystals diffracted to high resolution and crystal structure of CTSL-YLAL mutant happens to be the highest resolution structure of pro-form of CTSL till date. High resolution crystal structures enable us to identify the structural basis for these altered activities of the protease mutants.

Our results show that with careful selection of mutations in the S2 pocket of human cathepsin L, a mutant of CTSL can be generated that gain proline specificity for peptide substrate having proline at P2 position. This proline-specific mutant though can cleave peptide substrates with Pro in P2 position; it is incapable of cleaving intact triple helical collagen unless an appropriate GAG-binding surface is provided. This can be done by replacing only three surface residues of CTSL, which results in a collagenolytic human cathepsin L. Our earlier studies on recombinant human cathepsin K showed that disease-state condition in autosomal recessive genetic disease pycnodysostosis is not always due to loss of function of CTSK mutant, in some cases other factors associated with conformational changes may lead to the diseased condition (**42**). In that study, we have identified some disease related mutations of human cathepsin K where protease activity and specificities are altered without perturbing GAG dependent collagenase activity. Since the recombinant human Cathepsin L collagenase generated in the present study was prepared using a human enzyme template, it may have inclusive applicability for various pharmacological and medical applications due to its expected better immuno-compatibility. While reiterating the requirement for GAG-binding for efficient degradation of triple helical collagen by cysteine cathepsins together with proline-specificity, our results also pave the way for designing proteins with desirable enzymatic properties.

## Materials & Methods

### Construction of Proline-Specific Pro-CTSL and Generation of GAG-Binding Site on Proline-Specific Pro-CTSL

A Pro-CTSL clone was generated by sub-cloning the human Pre-Pro-CTSL ORF clone (in pMD18-T Simple Vector) obtained from Sino Biologicals Inc., Beijing, China, in pET30 Ek/LIC expression vector (Novagen, USA) using appropriate primers so as to delete the first 17 amino acids comprising the Pre- or the Signal peptide sequence. This wild-type Pro-CTSL clone was used as template for the subsequent mutagenesis studies. The Pro-CTSL protein expressed as inclusion bodies, which were solubilized with urea, purified by Ni-NTA affinity chromatography under denaturing conditions and refolded by dilution method as previously standardized in our laboratory (44; 45). Refolded proteins were concentrated at 4°C on a 200 ml Amicon stirred cell ultrafiltration unit using an YM 10 membrane with a 10 NMWL cut-off. Concentrations of the refolded proteins were determined from Bradford dye binding assay (46) and from the absorbance at 280 nm, using thecalculated extinction coefficient for the total translated product (pro, mature and vector-specific tagging sequence) (47). The refolded proteins were analyzed by SDS–PAGE for purity.

The cDNA of wild-type Pro-CTSL clone described above was used for introducing the mutations. SDM was carried out with appropriate primers (Table 4) using QuikChange^®^ Site-Directed Mutagenesis Kit (Stratagene, La Jolla, CA, USA) according to manufacturer’s instruction.

**Table 4:**
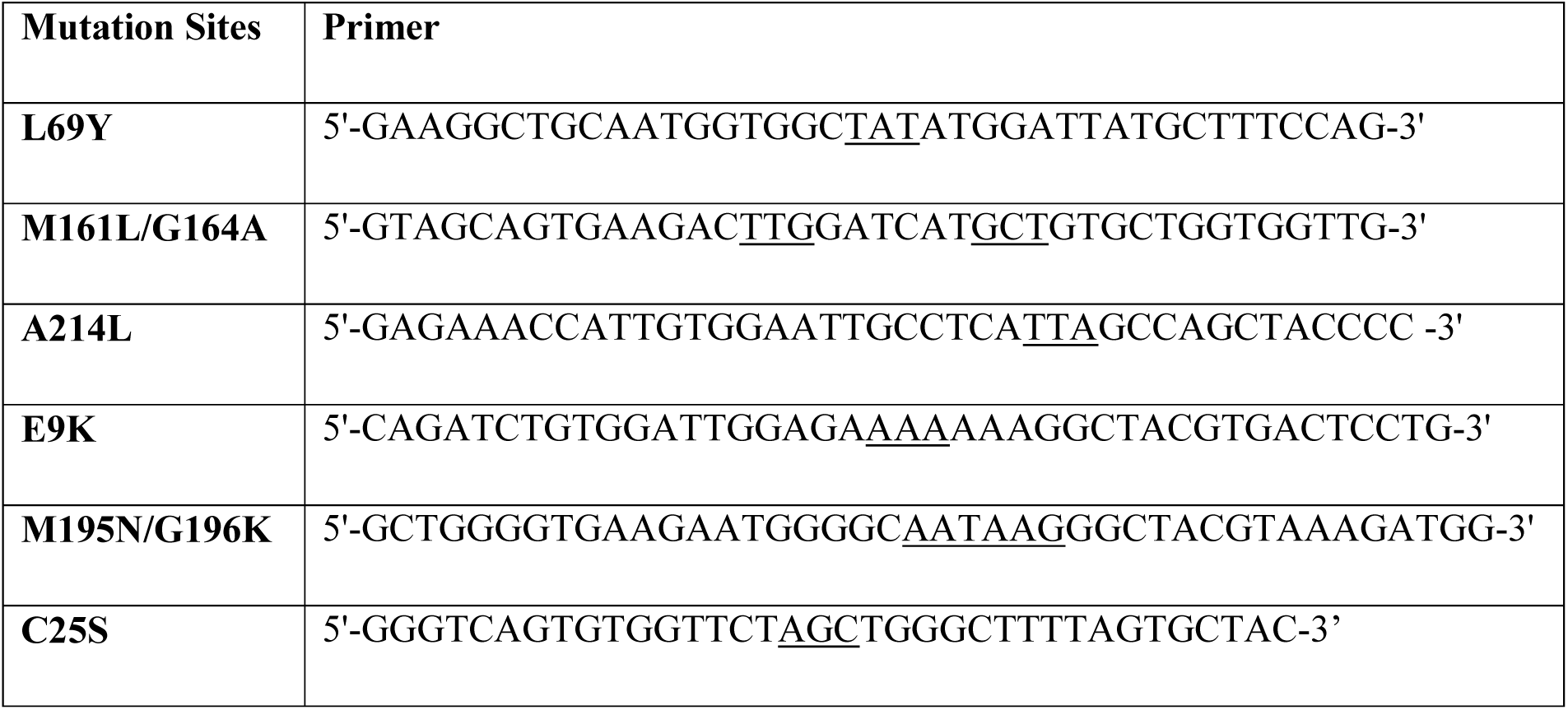
Mutagenic oligonucleotides used for site-directed mutagenesis (mature CTSL numbering)

**Table 5.** Data collection and refinement statistics.

|  | CTSL-YLAL | CTSL-C4S |
| --- | --- | --- |
| **Data statistics:** |  |  |
| Beam line | BM14, ESRF, Grenoble | BM14, ESRF, Grenoble |
| Wavelength (Å) | 1.0 | 1.0 |
| Resolution range | 50.0-1.46 (1.49- 1.46) | 50.0 - 1.80 (1.83 - 1.80) |
| Space group | P 21 21 21 | P 21 21 21 |
| Unit cell (Å) | 42.22 67.58 103.24 | 63.84 64.17 92.86 |
| Total reflections | 314047 | 161264 |
| Unique reflections | 49550 (1641) | 34526 (1491) |
| Multiplicity | 6.3(4.7) | 4.7(4.3) |
| Completeness (%) | 94.4 (63.6) | 96.8 (85.4) |
| Mean I/sigma(I) | 45.3(3.53) | 22.3(1.44) |
| Wilson B-factor | 17.32 | 24.84 |
| R-merge | 0.084(0.501) | 0.036(0.507) |
| R-meas | 0.092(0.427) | 0.063(0.709) |
| R-pim | 0.035(0.180) | 0.028(0.319) |
| **Refinement statistics:** |  |  |
| Resolution range used in refinement | 25.56 - 1.46 (1.51 - 1.46) | 32.41 - 1.80 (1.87 - 1.80) |
| Reflections used in refinement | 49439 (3490) | 34468 (2945) |
| Reflections used for R-free | 1998 (140) | 2016 (179) |
| R-work | 0.1916 (0.2726) | 0.1713 (0.2953) |
| R-free | 0.2380 (0.3192) | 0.2148 (0.3365) |
| Number of non-hydrogen atoms | 2879 | 2942 |
| macromolecules | 2586 | 2537 |
| ligands | 63 | 249 |
| solvent | 230 | 156 |
| Protein residues | 314 | 314 |
| RMS(bonds) | 0.032 | 0.010 |
| RMS(angles) | 1.57 | 0.99 |
| Ramachandran favored (%) | 95.45 | 96.43 |
| Ramachandran allowed (%) | 3.90 | 2.92 |
| Ramachandran outliers (%) | 0.65 | 0.65 |
| Rotamer outliers (%) | 5.38 | 2.23 |
| Clash score | 15.30 | 10.39 |
| Average B-factor | 28.66 | 34.33 |
| macromolecules | 27.83 | 31.84 |
| ligands | 46.36 | 57.69 |
| solvent | 33.15 | 37.63 |
| PDB code: | 6JD8 | 6JD0 |

Statistics for the highest-resolution shell are shown in parentheses.
R-merge = $\frac{\sum_h \sum_i |I_{h,i} - I_h|}{\sum_h \sum_i I_{h,i}}$ , where $I_h$ is the mean intensity of $i$ observations of symmetry related reflections of $h$ .
$R_{pim} = \frac{\sum_{hkl} [n/(n-1)]^{1/2} \sum_i |I_i(hkl) - \langle I(hkl) \rangle|}{\sum_{hkl} \sum_i I_i(hkl)}$ , where $n$ is the multiplicity, other variables as defined for R-merge.
R-work = $\sum |F_o - F_c| / \sum F_o$ . R-free is the cross-validation R factor for the test set (5%) of reflections omitted in model refinement.

A quadruple mutant (L69**Y**/M161**L**/G164**A**/A214**L**) of pro-CTSL (mature CTSL numbering) was generated to create a proline-specific CTSL (named CTSL-**YLAL**). GAG (Chondroitin 4-sulfate, C 4-S) binding region was generated on the Proline-specific pro-CTSL, CTSL-YLAL, by mutating three relevant amino acids (Y9K; M195N; G196K, mature CTSL numbering) on the surface of the protein taking signatures from Cathepsin K. This mutant was named CTSL-C4S mutant. The designated mutations in each variant were verified by automated DNA sequencing using the MegaBACE™ 1000 sequencing system (Amersham Biosciences, USA). All the mutants were over-expressed, purified and refolded as for wild type pro-CTSL with minor changes.

For structural studies, the active site Cysteine (C25) was replaced with Serine in both, CTSL-YLAL and CTSL-C4S mutant to create non-active proteases. Refolded CTSL-YLAL-C25S and CTSL-C4S-C25S proteins were concentrated and purified further by gel-filtration chromatography on a Sephacryl S-100 (Sigma-Aldrich, USA) column that had been pre-equilibrated with 20 mM Tris–HCl pH 8.0, 300 mM NaCl and 1% glycerol. The protein was eluted in the same buffer and the peak fractions were collected and concentrated to 10–15 mg/ml in same column buffer. Purity of the proteins was checked by 15% SDS–PAGE analysis.

### Substrate Gel Zymography and Activation of Precursor Proteins

Proper refolding of the recombinant active proteases, wild type (pro-CTSL) and mutants (CTSL-YLAL & CTSL-C4S mutant), from inclusion bodies was initially screened by Substrate Gel Zymography using 0.1% gelatin as a substrate using a protocol described previously (**48**). Briefly, the activated pro-enzymes were diluted with 4X non-reducing SDS-PAGE sample buffer, incubated for 20 min at 37°C, and separated by SDS-PAGE on 12% gels containing 0.1% gelatin at 4°C. After electrophoresis, the gels were washed with 2.5% Triton X-100 in 20 mM Tris/HCl, pH 8.0 for 30 min, and subsequently washed extensively with distilled water to remove the detergent completely. The gel was then incubated in Activation buffer (100 mM Na-acetate, pH 5.5, 1mM EDTA, 2 mM DTT) at 37°C for 30 min. Following this, the gel was incubated overnight at 37°C in the Activation buffer without DTT protein digestion, and then stained with Coomassie Brilliant Blue R-250. Proteinase activity appears as clear bands on a dark blue background.

To obtain active mature proteases, the activation protocol was optimized initially with the recombinant wild type Pro-CTSL by incubating the protease precursor at various temperatures ranging from 30 °C to 50 °C for varying time periods in presence or absence of activators like Cysteine and DTT in buffers with pH ranging from 3.5 to 6.5. The zymogen activation process was followed by assaying the enzyme activity with pEFLNA at 410 nm at the different stages of activation. Subsequently, an optimized protocol whereby the protease precursors are incubated in 100 mM sodium acetate buffer pH 4.0 containing 2 mM EDTA at 45 °C for 10 min in presence of 20 mM Cysteine was used for activation of the wild type as well as all the mutant zymogens to mature form.

### Substrate Assays Using p-Nitroanilide Peptides

Steady state kinetics for the wild type and the mutant proteases was determined with synthetic chromogenic substrates containing p-nitroanilide, pE**F**L↓NA, V**L**K↓-pNA and G**P**R↓-pNA (Sigma Aldrich, USA). The P2 residue and cleavage sites are shown in bold and arrow respectively. The pro-enzymes were activated at low pH as described in previous section and immediately added to a pre-determined amount of peptidyl substrates in 50 mM sodium acetate buffer pH 5.5 containing 2 mM EDTA, 2 mM cysteine and the enzyme activity was followed by monitoring the release of p-nitroaniline (pNA) from the substrates by measuring absorbance at 410 nm using an extinction coefficient of 8800 M^-1^ cm^-1^ for p-NA (**49**), in a UV/Vis spectrophotometer (Nicolet Evolution 100; Thermo Electron Corporation, Rockville, MD, USA). *K*_M_ and *V*_max_ values were determined by nonlinear regression analysis by fitting the reaction velocities into the Michaelis–Menten equation curve using the software Graphpad PRISM version 6 (http://www.graphpad.co/prism). The *k*_cat_ values were obtained using the equation *k*_cat_ = *V*_max_/[E]_T_, where [E]_T_ is the total concentration of the active enzyme, the values of which were measured by active-site titration with the irreversible inhibitor E-64 (trans-Epoxysuccinyl-L-leucylamido(4-guanidino)butane, which binds in a 1:1 molecular ratio to the active sites of most cysteine proteases (**50**).

### Collagen degradation assays and GAG Binding Studies

Collagen degradation assays were performed in absence and presence of GAG (chondroitin 4-sulfate) for the wild type, proline-specific CTSL-YLAL mutant and the CTSL-C4S mutant of CTSL. The pro-enzymes were first activated to mature form as indicated above. After activation, collagen degradation assays were performed in 0.1 M sodium acetate buffer, pH 6.5, containing 1 mM EDTA. In all experiments, soluble type I collagen from calf skin and rabbit skin collagen were used as the substrate at a concentration of 0.4 mg mL^-1^. Collagen digestion was performed at 28 °C and 37 °C for 2h, 4h and overnight in the presence and absence of 0.15% (w/v) C4-S. The digestion reaction was stopped by the addition of 10 μM of E-64. Aliquots were withdrawn at regular time intervals and analyzed by SDS/PAGE on 7.5% polyacrylamide gels. In all experiments, the BioRad Dual colour marker was used for molecular weight calibration.

The effect of GAG on the kinetic parameters was studied by measuring the enzymatic activity of the CTSL-C4S mutant in presence and absence of C4-S and heparin. The assay protocol was same as described earlier except that after activation of the zymogen, 0.15 % GAG was added just before addition of the peptidyl substrates.

Also, different GAGs like chondroitin 4-sulfate, heparin sulphate and heparin were used to study their binding effects in electrophoretic mobility of CTSL-YLAL and CTSL-C4S mutants in native gel. After incubation with a particular GAG, the proteins were analysed by Native-PAGE where a 10% polyacrylamide gel was prepared without SDS and the samples were incubated with a 2X sample buffer without SDS for 10 minutes at room temperature. The gels were run at 4 °C in a Tris-glycine buffer system again without SDS so as to avoid any damage to the proteins incubated with respective GAGs.

### Crystallization, data Collection and structure determination

Purified pro-form of the inactive mutant proteins, CTSL-YLAL-C25S and CTSL-C4S-C25S, from gel filtration was concentrated to 10-15 mg/mL and initial crystallization screening was set up using hanging drop vapour diffusion method with Structure Screen 1 and 2 (MD1-01 & MD1-02) of Molecular Dimensions Limited (United Kingdom). Diffraction quality crystals appeared within one week with 20% PEG 4000 in 100 mM Na-Hepes (pH 7.5) (Condition 31 of MD1-01)at 20°C. The crystals were cryo-protected with the above precipitant containing 15–30% glycerol and flash-frozen in liquid nitrogen for transportation for data collection.

X-ray diffraction data-sets were collected from flashed frozen crystals at 100K on a Mar-CCD detector at BM14 beam-line ESRF, Grenoble, France. The diffraction data were processed and scaled using HKL2000 program (51). Molecular replacement were carried out using PHASER (**52**) in PHENIX with the pro-cathepsin L wild-type structure (pdb id: 1CJL) as a model. For CTSL-YLAL-C25S and CTSL-C4S-C25S mutants, the mutated residues have been replaced by Ala residues in the model. These residues were placed according to electron density guideline. Iterative rounds of model building with Coot (**53**) and refinement using PHENIX (54) and REFMAC5 (55) were performed using a translation-liberation-screw model of the atomic displacement parameters. We could identify different ligand molecules and ions in well defined electron density maps in accordance with the buffers used during refolding, purification and crystallization and alternate conformations of different amino acid side-chains and some water molecules were placed in the final stage of refinement. Final model of the asymmetric unit of CTSL-YLAL mutant consists of a pro-CTSL-YLAL molecule, 225 water molecules (with 5 alternate conformations) and 16 different ligand molecules (ethylene glycol, glycerol, ethanol etc) and those of CTSL-C4S mutant are a pro-CTSL-C4S molecule, 156 water molecules and 35 different ligand molecules (ethylene glycol, glycerol, propanol, ethanol etc) and ion molecules (sulphate, chloride and sodium) (Table 5). Co-ordinates and structure factors were deposited in protein data bank (PDB) with accession codes 6JD8 and 6JD0 for CTSL-YLAL and CTSL-C4S mutants respectively. Refinement and diffraction data statistics are summarized in Table 5.

## Acknowledgements

The authors are grateful to Dr. Deepak Nair of RCB, Faridabad, India for help in Syncrotron data collection at ESRF, France, through DBT-ESRF collaborative project. This work was supported by Department of Atomic Energy (MSACR project to SINP) and WOS ‘A’ scheme, Department of Science and Technology (SR/WOS-A/LS-468/2012), Government of India.

## Author contributions

DC and SB conceived the project, DC performed all biochemical and molecular biology work. SB performed crystal structure refinement and analyses work. SB wrote the paper with DC.

## REFERENCES

1. Alberts, B., Johnson, A., Lewis, J., Raff, M., and Roberts, K. (2002) Molecular Biology of the Cell. Garland Science: New York, NY, USA

2. van der Rest, M., and Garrone, R. (1991) Collagen family of proteins. FASEB J. 5, 2814–2823

3. Shoulders, M. D., and Raines, R. T. (2009) Collagen structure and stability. Annu Rev Biochem. 78, 929–958

4. Everts, V., Aronson, D. C., and Beertsen, W. (1985) Phagocytosis of bone collagen by osteoclasts in two cases of pycnodysostosis. Calcif. Tissue Int. 37, 25–31

5. Gelb, B. D., Shi, G. P., Chapman, H. A., and Desnick, R. J. (1996) Pycnodysostosis, a lysosomal disease caused by cathepsin K deficiency. Science 273, 1236–1238

6. Kiviranta, R., Morko, J., Alatalo, S. L., NicAmhlaoibh, R., Risteli, J., Laitala-Leinonen, T., and Vuorio, E. (2005) Impaired bone resorption in cathepsin K-deficient mice is partially compensated for by enhanced osteoclastogenesis and increased expression of other proteases via an increased RANKL/OPG ratio. Bone.;36, 159–72

7. Murphy, G., and Hembry, R. M., (1992) Proteinases in rheumatoid arthritis. Journal of Rheumatology. 19(32), 61–4

8. Galis, Z. S., Sukhova, G. K., Kranzhöfer, R., Clark, S., and Libby, P. (1995) Macrophage foam cells from experimental atheroma constitutively produce matrix-degrading proteinases. Proceedings of the National Academy of Sciences USA. 92(2), 402–6

9. Hautamaki, R. D., Kobayashi, D. K., Senior, R. M.,and Shapiro, S. D. (1997) Requirement for macrophage elastase for cigarette smoke-induced emphysema in mice. Science. 277(5334), 2002–4

10. Werb, Z., Vu, T. H., Rinkenberger, J. L., and Coussens, L. M. (1999) Matrix-degrading proteases and angiogenesis during development and tumor formation. APMIS. 107(1-6), 11–18

11. Goshev, I., Gousterova, A., Vasileva-Tonkova, E., and Nedkov, P. (2005) Characterization of the enzyme complexes produced by two newly isolated thermophylic actinomycete strains during growth on collagen-rich materials. Process Biochem. 40, 1627–1631

12. Kanth, S. V., Venba, R., Madhan, B., Chandrababu, N. K., and Sadulla, S., (2008) Studies on the influence of bacterial collagenase in leather dyeing. Dye Pigment. 76, 338–347

13. Agren, M. S., Taplin, C. J., Woessner, J. F. Jr., Eagisteim, W. H., and Mertz, P. M., (1992) Collagenase in wound healing: effect of wound age and type. J Invest Dermatol. 99, 709–714

14. Püllen, R., Popp, R., Volkers, P., and Füsgen, I., (2002) Prospective randomized double-blind study of the wound-debriding effects of collagenase and fibrinolysin/deoxyribonuclease in pressure ulcers. Age Ageing. 31, 126–130

15. Shmoilov, A. M., Rudenskaya, G. N., Isev, V. A., Baydakov, A.V., and Zhantiev, R. D., (2006) A comparative study of collagenase complex and new homogeneous collagenase preparations for scar treatment. J Drug Deliv Sci Technol. 16, 285–292

16. Lima, C. A., Campos, J. F., Filho, J. L., Converti, A., da Cunha, M. G., and Porto, A. L. (2014) Antimicrobial and radical scavenging properties of bovine collagen hydrolysates produced by Penicillium aurantiogriseum URM 4622 collagenase. J Food Sci Technol. 52(7), 4459–4466

17. Chung, L., Shimokawa, K. I., and Nagase, H. (2000). Structural requirements for collagenolytic activity of matrix metalloproteinase 1 (MMP-1). Kluwer Academic Publishers, Dordrecht, The Netherlands

18. Chung, L., Dinakarpandian, D., Yoshida, N., Lauer-Fields, J. L., Fields, G. B., Visse, R., and Nagase, H. (2004) Collagenase unwinds triple-helical collagen prior to peptide bond hydrolysis. EMBO J. 23, 3020–3030

19. Etherington, D. J. (1972) The nature of the collagenolytic cathepsin of rat liver and its distribution in other rat tissues. Biochem J. 127, 685–692

20. Etherington, D. J.,and Evans, P. J. (1977) The action of cathepsin B and collagenolytic cathepsin in the degradation of collagen. Acta Biol Med Ger. 36, 1555–1563

21. Garnero, P., Borel, O., Byrjalsen, I., Ferreras, M., Drake, F. H., McQueney, M. S., Foged, N.T., Delmas, P. D., and Delaisse, J. M. (1998) The collagenolytic activity of cathepsin K is unique among mammalian proteinases. J Biol Chem. 273, 32347–32352

22. Kafienah, W., Bromme, D., Buttle, D. J., Croucher, L. J., and Hollander, A. P. (1998) Human cathepsin K cleaves native type I and II collagens at the N-terminal end of the triple helix. Biochem J. 331, 727–732

23. Li, Z., Hou, W. S., and Bromme, D. (2000) Collagenolytic activity of cathepsin K is specifically modulated by cartilage-resident chondroitin sulfates. Biochemistry. 39, 529–536

24. Everts, V., Korper, W., Hoeben, K. A., Jansen, I. D., Bromme, D., Cleutjens, K. B., Heeneman, S., Peters, C., Reinheckel, T., Saftig, P., and Beertsen, W. (2006) Osteoclastic bone degradation and the role of different cysteine proteinases and matrix metalloproteinases: differences between calvaria and long bone. J Bone Miner Res. 21(9), 1399–408

25. Georges, S., Ruiz, Velasco C., Trichet, V., Fortun, Y., Heymann, D., and Padrines, M. (2009) Proteases and bone remodelling. Cytokine Growth Factor Rev. 20(1), 29–41

26. Charni-Ben Tabassi, N., Desmarais, S., Bay-Jensen, A. C., Delaisse, J. M., Percival, M. D., and Garnero, P. (2008) The type II collagen fragments Helix-II and CTX-II reveal different enzymatic pathways of human cartilage collagen degradation. Osteoarthr Cartil. 16, 1183–91

27. Hill, P. A., Buttle, D. J., Jones, S. J., Boyde, A., Murata, M., Reynolds, J. J., and Meikle, M. C. (1994) Inhibition of bone resorption by selective inactivators of cysteine proteinases. J Cell Biochem. 56, 118–30

28. Bromme, D. and Kaleta, J. (2002) Thiol-dependent cathepsins: pathophysiological implications and recent advances in inhibitor design. Curr. Pharm. Des. 8, 1639–1658

29. Gottesman, M.M. and Smith, S.M. (1989) Activity and deletion analysis of recombinant human cathepsin L expressed in Escherichia coli. J. Biol. Chem. 264, 20487–20495

30. Nägler, D.K. and Menard, R. (2003). Family C1 cysteine proteases: Biological diversity or redundancy? Biol. Chem. 384, 837–843

31. Li, Z., Hou, W. S., Escalante-Torres, C. R., Gelb, B. D., and Bromme, D. (2002). Collagenase activity of cathepsin K depends on complex formation with chondroitin sulfate, J. Biol. Chem. 277(32), 28669–28676

32. Borel, O., Gineyts, E., Bertholon, C., and Garnero, P. (2012) CathepsinK preferentially solubilizes matured bonematrix, Calcified Tissue International. 91(1), pp. 32–39

33. Li, Z., Kienetz, M., Cherney, M. M., James, M. N., and Brömme, D. (2008) The crystal and molecular structures of a cathepsin K: chondroitin sulfate complex. J. Mol. Biol. 383, 78–91

34. Yasuda, Y., Li, Z., Greenbaum, D., Bogyo, M., Weber, E., and Brömme, D. (2004). Cathepsin V, a novel and potent elastolytic activity expressed in activated macrophages. J. Biol. Chem. 279, 36761–36770

35. Schechter, I., and Berger, A. (1967) On the size of the active site in proteases. I. Papain. Biochem. Biophys. Res. Commun. 27, 157–162

36. Coulombe, R., Grochulski, P., Sivaraman, J., Ménard, R., Mort, J. S., and Cygler, M. (1996). Structure of human procathepsin L reveals the molecular basis of inhibition by the prosegment. The EMBO Journal. 15, 5492–5503

37. Lecaille, F., Vandier, C., Godat, E., Hervé-Grépinet, V., Brömme,, D., and Lalmanach, G. (2007). Modulation of hypotensive effects of kinins by cathepsin K. Arch. Biochem. Biophys. 459, 129–136

38. Li, Z., Yasuda, Y., Li, W., Bogyo, M., Katz, N., Gordon, R. E., Fields, G. B., and Brömme, D. (2004) Regulation of collagenase activities of human cathepsins by glycosaminoglycans, J. Biol. Chem. 279, 5470–5479

39. Nosaka, A. Y., Kanaori, K., Teno, N., Togame, H., Inaoka, T., Takai, M., and Kokubo, T. (1999) Conformational studies on the specific cleavage site of Type I collagen (alpha-1) fragment (157–192) by cathepsins K and L by proton NMR spectroscopy. Bioorg Med Chem. 7, 375–379

40. Ishidoh, K., and Kominami, E. (1995) Procathepsin L degrades extracellular matrix proteins in the presence of glycosaminoglycans in vitro. Biochem Biophys Res Commun. 217, 624–631

41. Cherney, M. M., Lecaille, F., Kienitz, M., Nallaseth, F. S., Li, Z., James, M. N., and Brömme, D. (2011) Structure-activity analysis of cathepsin K/chondroitin 4-sulfate interactions. J. Biol. Chem. 286, 8988–8998

42. Roy, S., Das Chakraborty, S., and Biswas, S. (2018) Not all pycnodysostosis-related mutants of human cathepsin K are inactive – crystal structure and biochemical studies of an active mutant I249T. FEBS Journal. 285, 4265–4280

43. Novinec, M., Kovacic, L., Lenarcic, B., and Baici, A. (2010) Conformational flexibility and allosteric regulation of cathepsin K Biochem. J. 429, 379–389

44. Aich, P., and Biswas, S. (2018) Highly Conserved Arg Residue of ERFNIN Motif of Pro-Domain is Important for pH-Induced Zymogen Activation Process in Cysteine Cathepsins K and L. Cell Biochem Biophys. 76, 219–229

45. Alam, B., and Biswas, S. (2019) Inhibition of *Plasmodium falciparum* cysteine protease falcipain-2 by a human cross-class inhibitor serpinB3: A mechanistic insight BBA - Proteins and Proteomics. 1867, 854–865

46. Bradford, M. M. (1976) A rapid and sensitive method for the quantitation of microgram quantities of protein utilizing the principle of protein-dye binding. Anal. Biochem. 72, 248–254

47. Gill, S. C. and von Hippel, P. H. (1989) Calculation of protein extinction coefficients from amino acid sequence data. Anal. Biochem. 182, 319–326

48. Choudhury, D., Biswas, S., Roy, S., and Dattagupta, J. K. (2010) Improving thermostability of papain through structure-based protein engineering. Protein Eng Des Sel. 23, 457–467

49. Mole, J. E., and Horton, H. R. (1973) Kinetics of papain-catalyzed hydrolysis of N-benzoyl-L-arginine-p-nitroanilide. Biochemistry. 12, 816–22

50. Barrett, A. J., Kembhavi, A. A., Brown, M. A., Kirschke, H., Knight, C. G., Tamai, M., and Hanada, K. (1982) L-trans-Epoxysuccinyl-leucylamido(4-guanidino)butane (E-64) and its analogues as inhibitors of cysteine proteinases including cathepsins B, H and L. Biochem J. 201, 189–198

51. Otwinowski, Z., and Minor, W. (1997) Processing of X-ray diffraction data collected in oscillation mode. Methods Enzymol. 276, 307–326

52. McCoy, A. J., Grosse-Kunstleve, R. W., Adams, P. D., Winn, M. D., Storoni, L. C., andRead, R. J. (2007) Phaser: crystallographicsoftware. J Appl Crystallogr. 40, 658–674. PMID: 9461840

53. Emsley, P., and Cowtan, K. (2004) COOT: model-building tools for molecular graphics. Acta Crystallogr D Biol Crystallogr. 60, 2126–2132

54. Adams, P. D., Grosse-Kunstleve, R. W., Hung, L. W., Ioerger, T. R., McCoy, A. J., Moriarty, N. W., Read, R. J., Sacchettini, J. C., Sauter, N. K., and Terwilliger, T. C. (2002) PHENIX:building new software for automated crystallographic structure determination. Acta Cryst Section D Biol Cryst. 58, 1948–1954

55. Murshudov, G. N., Vagin, A. A., andDodson, E. J. (1997) Refinement of macromolecular structures by the maximum-likelihood method. Acta Cryst D Biol Cryst. 53, 240–255

